# Probucol mitigates high-fat diet-induced cognitive and social impairments through disruption of redox-inflammation association

**DOI:** 10.1101/2023.09.05.556289

**Authors:** Han-Ming Wu, Na-Jun Huang, Yang Vivian Yang, Li-Ping Fan, Tian-Yu Tang, Lin Liu, Yue Xu, Dong-Tai Liu, Ze-Xin Cai, Xin-Yi Ren, Zheng-Hao Yao, Jian-Zhen Chen, Xi Huang, Cixiong Zhang, Xiang You, Chen Wang, Ying He, Zhi-Yun Ye, Wei Hong, Sheng-Cai Lin, Yi-Hong Zhan, Shu-Yong Lin

## Abstract

Obesity and its detrimental metabolic consequences are commonly recognized as risk factors for impairments in the central nervous system (CNS). However, the direct link between metabolic abnormalities and brain functions during high-fat feeding remains unclear. Here, we show that treatment with probucol, a cholesterol-lowering drug, counteracts the cognitive and social impairments induced by a high-fat diet in mice, while having no effect on mood disorders. Unexpectedly, the beneficial effects of probucol do not result from rectifying obesity or restoring glucose and lipid homeostasis, as evidenced by the lack of change in body weight, blood glucose and serum cholesterol levels. Interestingly, high-fat feeding led to association among the levels of redox factors, including oxidized low-density lipoprotein, glutathione and malondialdehyde, as well as a significant negative correlation between malondialdehyde levels and behavioral performance. Probucol treatment interrupts these linkages and differentially regulates the proteins for the generation of reactive oxygen species and reactive nitrogen species in the brain. These findings prompt a reconsideration of the mechanism of action of probucol, as well as the roles of altered metabolic profiles and free radicals in brain function.

## Introduction

Obesity and its negative metabolic consequences such as type 2 diabetes and hyperlipidemia, are generally regarded as conferring risk factors for impairments in the central nervous system (CNS) (1–5). Consumption of calorie-dense diets, which is considered one of the major factors contributing to the obesity pandemic, has been linked to cognitive dysfunction and mood disorders in humans (6–12). Numerous animal studies also show that high-fat diet (HFD) feeding that is routinely used to establish obese animal models leads to impaired learning and memory, and induces anxiety- and depression-like behaviors (13–19). Insulin resistance, altered lipid homeostasis, increased systemic oxidative stress and chronic inflammation, as well as dysfunctional vascularization under obese conditions intersect to promote the development of structural and molecular changes in the brain (20–25).

Various strategies, such as nutrition, anti-obesity drugs and exercise, have been tested to assess their effects in mitigating cognitive and mood impairments induced by HFD (26–29). As anticipated, most of these treatments demonstrated improvement in obesity-related metabolic abnormalities, including reductions in body weight, fat content, serum glucose levels, and insulin resistance to varying extents, alongside their diverse impacts on CNS. However, several studies indicate that the beneficial effects on cognitive decline and anxiety can be achieved without directly targeting the systemic metabolic changes induced by HFD (30–36). These findings suggest the presence of multiple pathways that mediate the impact of dietary intake on specific CNS abnormalities.

Redox homeostasis represents a prominent focus of investigation within these pathways. Lipid oxidation products, such as oxidized low-density lipoprotein (oxLDL) and malondialdehyde (MDA), as well as nitric oxide (NO) derived from the inducible nitric oxide synthase (iNOS), are positively correlated with obesity severity (37–40). These molecules play critical roles in neuroinflammation, which triggers the production of proinflammatory cytokines, including TNFα. Moreover, certain radical species directly interact with cellular macromolecules like lipids, DNA, and proteins. Consequently, these effects contribute to cellular component damage and tissue destruction (41–43). Nonetheless, it is evident that reactive oxygen species (ROS) also serve important roles in various physiological processes, such as brain development and plasticity, post-trauma angiogenesis, and the elimination of dysfunctional cells (44–46). In line with this perspective, although several small-scale clinical or laboratory studies demonstrate the beneficial effects of antioxidants in mitigating cognitive decline and mood disorders, (47–50), large-scale clinical trials generally do not yield significant positive outcomes (51, 52). Hence, modulating the activity and specificity of the antioxidants may be crucial for achieving context-dependent oxidant-antioxidant balance.

Probucol, known for its use as a cholesterol-lowering drug, has also demonstrated antioxidant properties by inhibiting the oxidative modification of low-density lipoprotein (LDL) (53, 54). The beneficial effects of probucol on cognitive dysfunction has been reported in pathological models, including mice injected with Amyloid beta proteins (55), those with D-galactose-induced cognitive deficits (56, 57), and those with drug or high-cholesterol diet-induced diabetes (58, 59). However, it remains unclear whether the cholesterol-reducing and antioxidant effects of probucol universally underlie these positive effects. Clinical trials suggest that the metabolic and antioxidative effects of probucol are relatively mild and primarily observed in individuals with very high cholesterol levels (60–63).

## Results

### Beneficial effects of probucol on cognitive abilities and social behavior in HFD-fed mice

To investigate the potential of probucol in mitigating the adverse effects of HFD on cognition and behavior, C57BL/6J mice were assigned to either a normal chow diet (NCD) or a 60% HFD for 8 weeks. Subsequently, the HFD-fed mice were divided into two groups based on their body weights. One group received 0.005% probucol (estimated to be 10-25 mg/kg/day) added to their drinking water, while the other two groups continued on the original diet without treatment. After 12 weeks of probucol intervention, the mice were subjected to the Morris water maze test. The untreated HFD-fed mice displayed reduced learning ability compared to the NCD-fed mice, evident by significantly increased latency to reach the hidden platform during the 7-day navigation task. In contrast, probucol treatment reduced the latency to a level comparable to that of the NCD-fed mice (Figure 1A). Following the navigation task, the mice underwent probe trials to evaluate their spatial memory. The untreated HFD-fed mice not only showed significantly increased latency to reach the platform area (Figure 1B) but also exhibited significantly reduced time spent in the target quadrant, fewer target-crossing events, and increased mean distance to the target (Figure 1, C and D). Conversely, these impairments were alleviated in the mice treated with probucol (Figure 1, B-D). Social behavior was also assessed using a three-chamber test. The probucol-treated mice displayed a significant preference for interacting with stranger mice over exploring an empty cage, whereas the untreated HFD-fed mice did not exhibit such a preference (Figure 1, E and F). These findings indicate that probucol treatment protects against HFD-induced damage to spatial learning, memory and social preference in mice.

**Figure 1.**
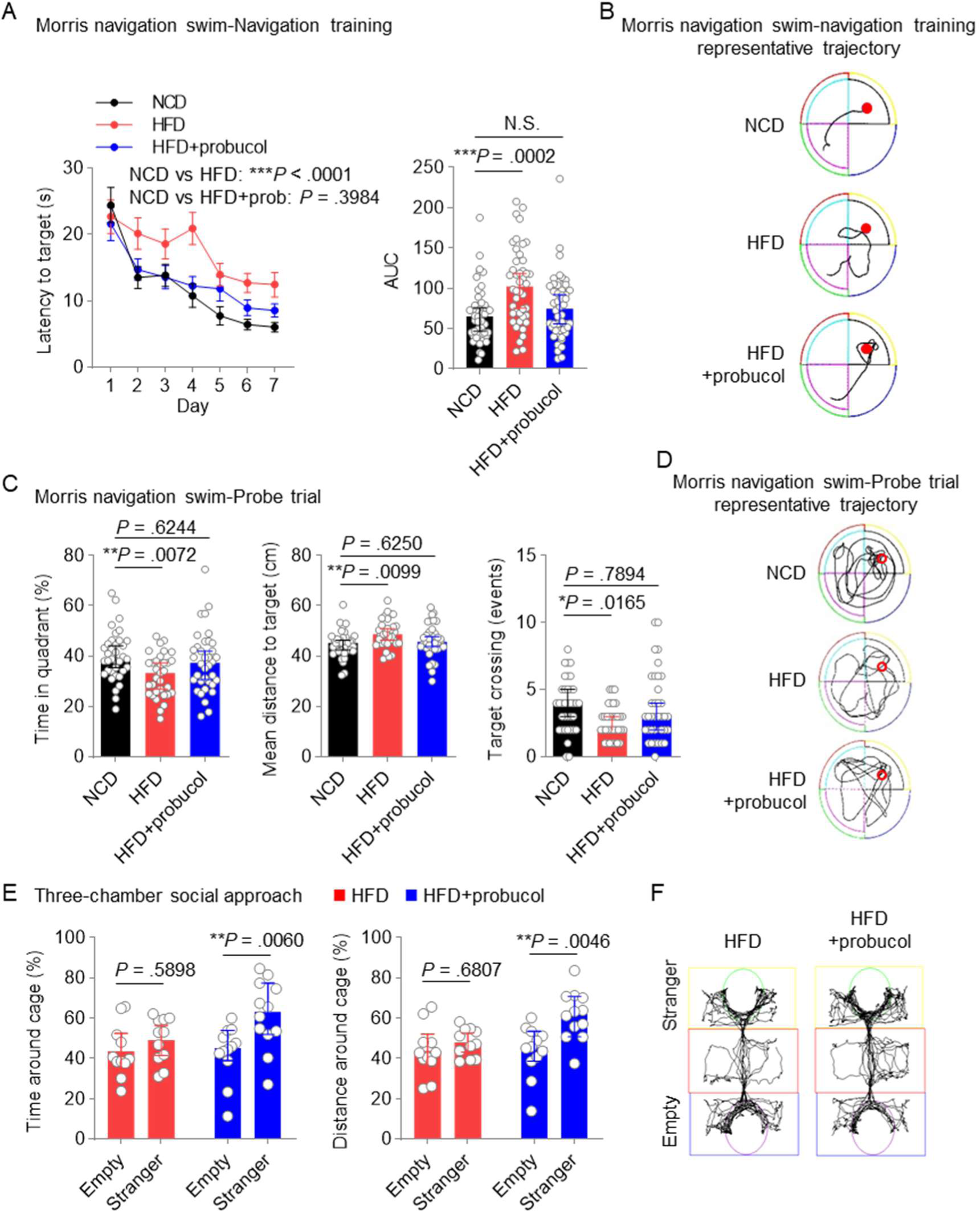
Probucol counteracts HFD-induced deficits in spatial and social cognition. (**A**) Performance of the mice in navigation training of the Morris swim navigation task. HFD-feeding started 8 weeks before treating the male C57BL/6J mice with 0.005% probucol in drinking water for 12 weeks. The training was stopped after more than 90% of the mice found the platform within 30 s for two consecutive days. Data of latency to target are shown in the left as mean ± SEM (n = 11 or 12 mice for each group, and 4 starting points per mice, ordinary two-way ANOVA followed by Tukey’s multiple comparisons test). The areas under the curve (AUC) of the latency to target of each mouse are shown in the right as individual values with median ± 95% CI (Kruskal-Wallis test followed by Dunn’s multiple comparisons test). (**B**) Representative partial trajectories of the mice in the probe trials showing latency to the platform. (**C**) Performance of the mice in the probe trials of the Morris swim navigation task. After reaching the standard of navigation training, the probe trials were conducted for mice described in **A** by removing the hidden platform. The percentages of the time in target quadrant, the mean distance to target, and target crossing numbers are shown as individual values with median ± 95% CI (n = 11 or 12 mice for each group, and 3 starting regions per mice, data of time in quadrant and mean distance to targets were compared by ordinary one-way ANOVA followed by Tukey’s multiple comparisons test, data of target crossing numbers were compared by Kruskal-Wallis test followed by Dunn’s multiple comparisons test). (**D**) Representative full trajectories of the mice in the probe trials of the Morris swim navigation task. (**E**) Social interaction of the mice in conventional three-chamber tasks. HFD-feeding started 8 weeks before treating the male C57BL/6J mice with 0.005% probucol in drinking water for 12 weeks. Stranger and Empty in the sociability task indicate the cage with a novel stranger male mouse and an empty cage, respectively. The percentages of the time spent around the cage and the percentages of distance around the cage are shown as individual values with median ± 95% CI (n = 11 or 12 mice for each group, two-way RM ANOVA). (**F**) Representative trajectory of the HFD-fed mice and probucol treated mice in conventional three-chamber tasks.

### Probucol has no effect on HFD-induced mood disorders in mice

It has been reported that HFD can also induce anxiety and depression-like behavior in mice. To determine whether probucol possesses anti-anxiety properties in HFD-fed mice, an elevated plus maze test was conducted. Both the untreated and probucol-treated HFD-fed mice exhibited significantly reduced downward exploration events, time spent, and distance traveled in the open arms compared to the NCD-fed mice (Figure 2A). These results suggest that anxiety levels were similarly developed in both groups of mice fed with HFD. Additionally, forced swim tests were conducted to assess the potential effects of probucol on depression-like behavior. Increased duration of immobility and reduced global activity were observed in both the untreated and probucol-treated HFD-fed mice compared to the NCD-fed mice (Figure 2B). These findings indicate that the beneficial effect of probucol are selective in targeting the cognitive performance affected by the HFD, but do not have an impact on mood disorders.

**Figure 2.**
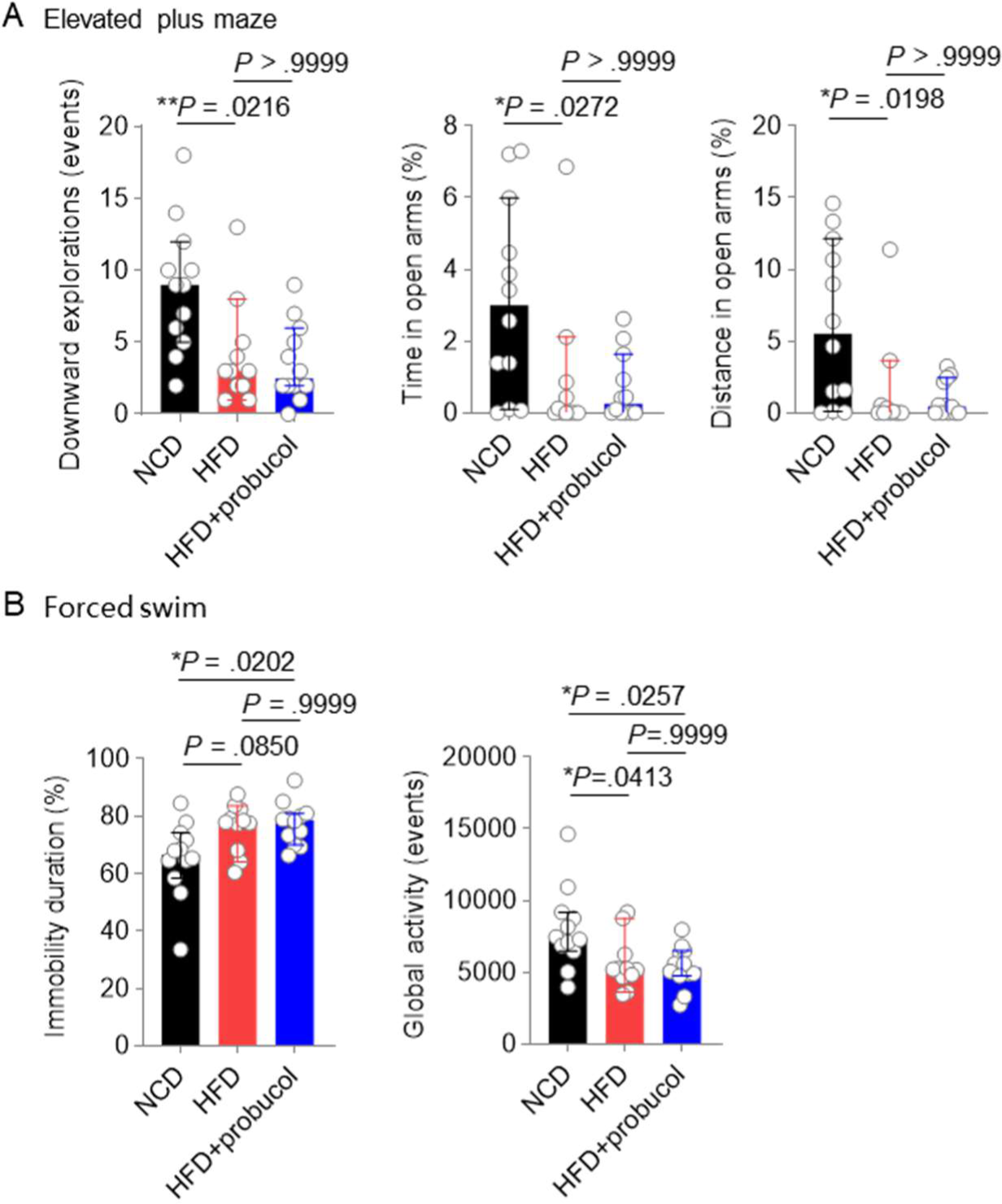
Probucol has no effect on anxiety and depression-like behaviors in HFD-fed mice. (**A**) Performance of the mice in the elevated plus maze. HFD-feeding started 8 weeks before treating the male C57BL/6J mice with 0.005% probucol in drinking water for 12 weeks. The data of downward explorations, time in open arms and distance in open arms are shown as individual values with median ± 95% CI (n = 11 or 12 mice for each group, Kruskal-Wallis test followed by Dunn’s multiple comparisons test). (**B**) Performance of the mice in the forced swim task. HFD-feeding started 8 weeks before treating the male C57BL/6J mice with 0.005% probucol in drinking water for 12 weeks. The immobility duration and global activity are shown as individual values with median ± 95% CI (n = 11 or 12 mice for each group, Kruskal-Wallis test followed by Dunn’s multiple comparisons test).

### Probucol selectively affects brain morphology without alleviating metabolic challenges from HFD

As dysregulated metabolism is considered a primary cause of various comorbidities associated with HFD, including impaired cognitive functions, we investigated the effects of probucol on the metabolic profiles of these mice. The HFD-fed mice showed significantly increases in body weight, blood glucose levels, total cholesterol and total LDL-cholesterol, which were not mitigated by probucol treatment (Figure 3, A-D). These results indicate that probucol does not have a noticeable effect on alleviating the global metabolic disorders induced by HFD, suggesting that its beneficial effects are not achieved through antagonizing the overall metabolic changes.

**Figure 3.**
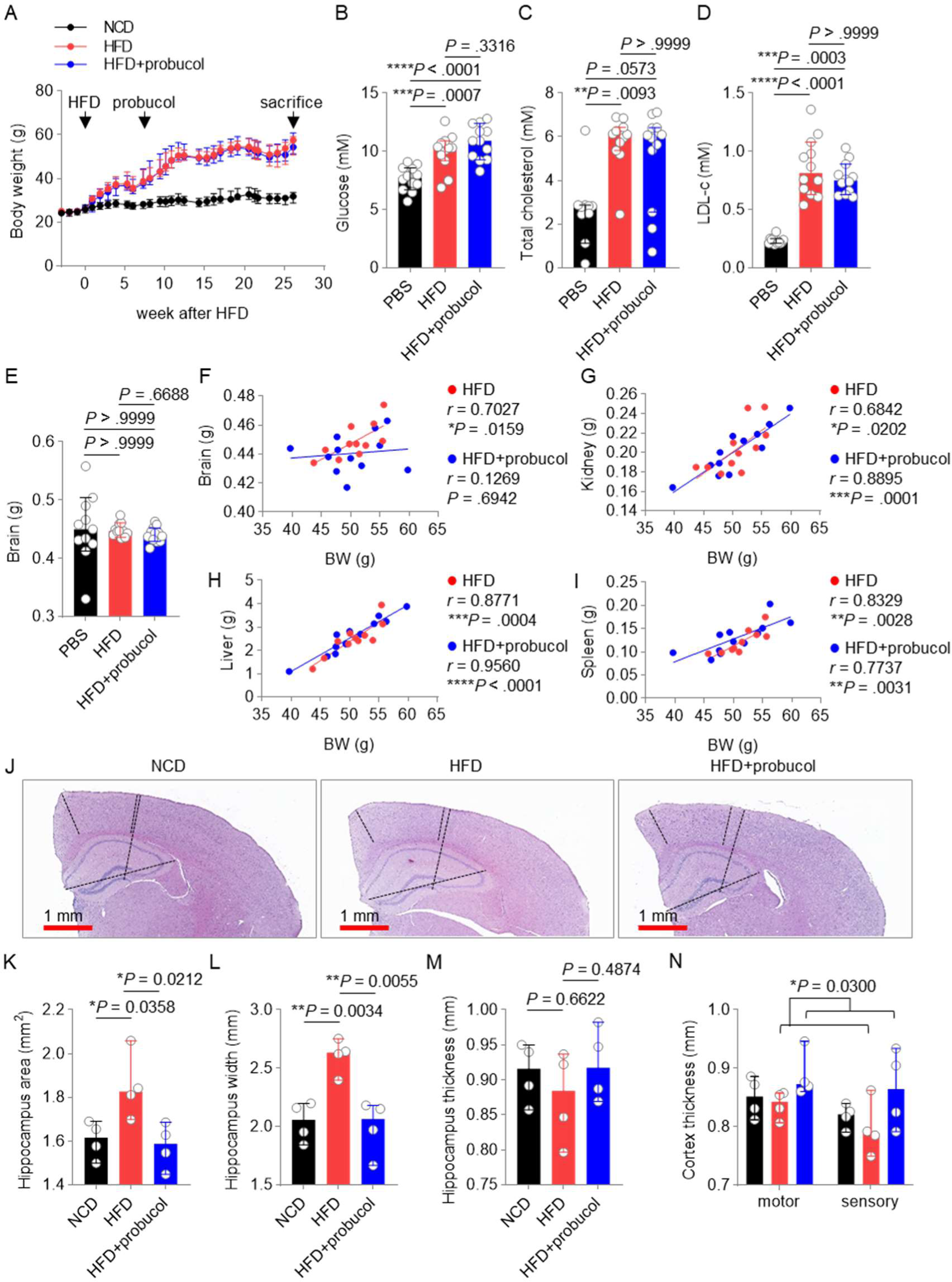
Probucol has no metabolic beneficial effects for HFD fed mice. (**A**) Probucol treatment has no effect on the body weight of the mice. Data are shown as median ± 95% confidence interval (CI). HFD-feeding started 8 weeks before treating the male C57BL/6J mice with 0.005% probucol in drinking water for 12 weeks. (**B-D**) Probucol treatment has no effect on blood glucose, serum total cholesterol and LDL-cholesterol of the mice. Blood of the mice were collected after 6-h fasting. Data are shown as individual values with median ± 95% CI (n = 12 mice for each group). The difference of the blood glucose between groups was compared by ordinary one-way ANOVA, followed by Tukey’s multiple comparisons test; those of total cholesterol and LDL-cholesterol were compared by Kruskal-Wallis test, followed by Dunn’s multiple comparison test. (**E**) Brain weights of the mice. Brains dissected from the mice were weighted and shown as individual values with median ± 95% CI (n = 11 or 12 mice for each group, Kruskal-Wallis test, followed by Dunn’s multiple comparisons test). (**F-I**) The correlations between body weights and the weights of organs such as the brain, kidney, liver and spleen of the HFD-fed mice and probucol treated mice. The degree of correlations is measured by Pearson r correlation. (**J-N**) Morphometric analysis of the effect of probucol on the mice brain. Hematoxylin and eosin (H&E) staining of the coronal sections containing the cerebral cortex and hippocampus of the mice (**J**). Data are shown as individual values with median ± 95% CI for the thickness of the cortex (**K**), the hippocampus area (**L**), the hippocampus width (**M**), and the thickness of hippocampus (**N**). The difference of the cortex thickness is compared by two-way ANOVA, followed by Tukey’s multiple comparisons test; those of the hippocampus parameters were compared by ordinary one-way ANOVA, followed by Tukey’s multiple comparisons test.

Reduced brain weight has been reported in mouse models of cognitive decline. We next determined the organ weights of these mice and found that neither HFD-feeding nor probucol treatment led to significant change in brain weight (Figure 3E). However, we observed a strong positive correlation between brain weight and body weight in the untreated HFD-fed mice, which was disrupted in the probucol-treated group (Figure 3F). In contrast, robust positive correlations between body weight and other organs such as the kidney, liver and spleen are consistently found in both the untreated and probucol-treated mice (Figure 3, G-I).

To further investigate the effects of HFD on brain regions vulnerable to cognitive decline, we performed quantitative morphometric measurements of the mice’s hippocampus and cortex (64), which are known to support flexible cognition and social behavior (65–67). We found that HFD-feeding significantly increased the total area and width of the hippocampus without changing its thickness. Probucol treatment restored the morphology of the hippocampus to a similar state as that of the NCD-fed mice (Figure 3, J-M). Additionally, mice treated with probucol showed increased thickness in the motor and sensory cortex compared to untreated HFD-fed mice, although the reduction in thickness was not statistically significantly in the untreated group (Figure 3N). These findings suggest that probucol selectively reshaped the connection between HFD-induced obesity and brain morphology, particularly in the hippocampus and cortex regions.

### Probucol’s influence on oxidative stress in HFD-fed mice

To explore the possibility of probucol alleviating HFD-induced oxidative stress, we conducted tests on the blood levels of oxidized low-density lipoprotein cholesterol (oxLDL-c), a major target of probucol. Surprisingly, we observed similar increases of oxLDL in both the untreated and probucol-treated HFD-fed mice compared to the NCD-fed mice (Figure 4A). Furthermore, the levels of malondialdehyde (MDA), an end product of lipid peroxidation and a major source of oxLDL-c modification, were even higher in the livers of probucol-treated mice (Figure 4B). These findings suggest that probucol does not function by antagonizing HFD-induced oxidative stress. We further analyzed the correlation between oxLDL-c and MDA levels. Consistent with previous reports, HFD-feeding established a positive correlation between oxLDL-c and MDA, which was dampened by probucol treatment (Figure 4C).

**Figure 4.**
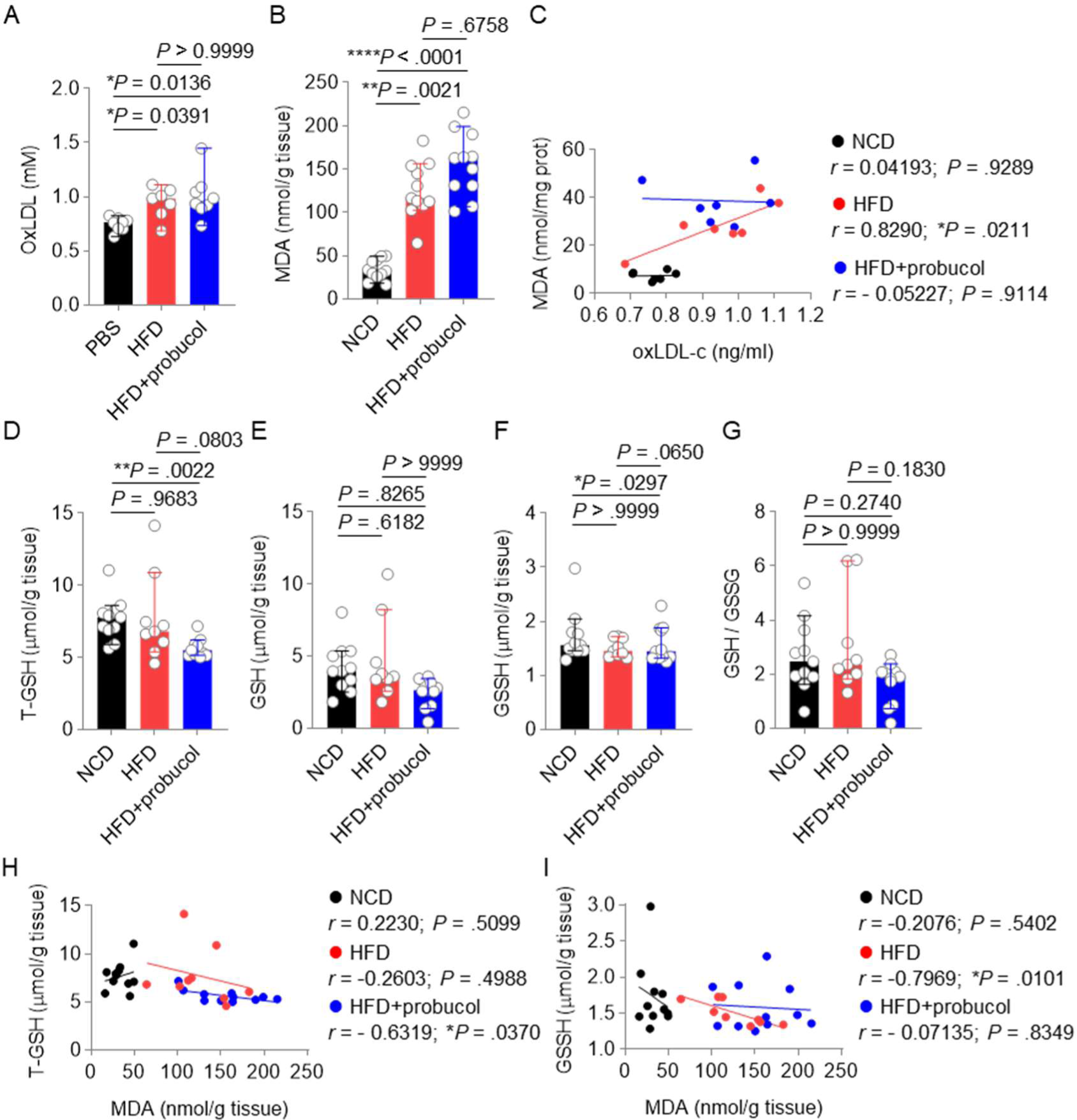
Impact of probucol feeding on mice redox status. (**A-C**) The levels of oxLDL-c (**A**) and MDA (**B**) of the mice. HFD-feeding started 8 weeks before treating the male C57BL/6J mice with 0.005% probucol in drinking water for 12 weeks. Data are shown as individual values with median ± 95% CI (n = 7 or 8 mice per group for oxLDL-c, n = 11 mice per group for MDA. Kruskal-Wallis test, followed by Dunn’s multiple comparisons test). The correlations between oxLDL-c and MDA are shown in (**C**). The degree of correlations is measured by Pearson r correlation. (**D-G**) The levels of T-GSH (**D**), GSH (**E**), GGSH (**F**), and the GSH:GSSH ratio (**G**) in the mice. Data are shown as individual values with median ± 95% CI (n = 9 or 11 mice for each group. Kruskal-Wallis test, followed by Dunn’s multiple comparisons test). (**H**, **I**) The impact of probucol treatment on the correlation between MDA and T-GSH o GSSH. The correlations between T-GSH and MDA (**H**), GGSH and MDA (**I**) are measured by Pearson r correlation.

We also investigated the levels of reduced glutathione (GSH), a crucial scavenger for reactive oxygen species (ROS), in these mice. Interestingly, probucol treatment significantly reduced total GSH levels compared to both the NCD-fed mice and the untreated HFD-fed mice. This reduction was achieved by decreasing the reductive GSH levels while leaving the oxidative GSSH levels unchanged. As a result, the GSH:GSSH ratio was also moderately reduced in probucol-treated mice (Figure 4, D-G). Consistently, probucol treatment established a negative correlation between total GSH and MDA, but not between GSSG levels and MDA (Figure 4, H and I). These findings indicate that probucol treatment disrupts the strong connection between MDA and the oxidization of LDL induced by HFD, while establishing a link between MDA and reductive GSH levels.

### Probucol’s protective effects on mice cognitive performance is associated with MDA levels

To investigated the association between the effect of probucol on MDA levels and its beneficial effects on cognition, we examined the correlations between MDA levels and performance in the Morris water maze test, as well as the forced swim test. We found that HFD-feeding established positive correlations between MDA levels and performance in the probe trial of the Morris water maze test (Figure 5A), indicating that MDA levels are specifically associated with cognitive performance rather than mood. Interestingly, oxLDL-c levels were found to be associated with better performance in the Morris water maze test in NCD-fed mice, but this association was reversed by HFD-feeding, resulting in poorer performance. However, probucol treatment disrupted this association. No significant correlations were observed between oxLDL-c and the depression-like behavior in any of the three groups of mice (Figure 5B). These findings suggest that spatial memory, but not depression-like behavior is associated with MDA and oxLDL-c levels. Furthermore, probucol may exert its effects by preventing the detrimental effects of oxidative products like MDA. Moreover, in alignment with the effect of probucol on brain weight, the positive correlation between brain weight and MDA levels induced by HFD was mitigated by probucol treatment, whereas no significant correlation was observed between brain weight and oxLDL-c levels (Figure 5, C and D).

**Figure 5.**
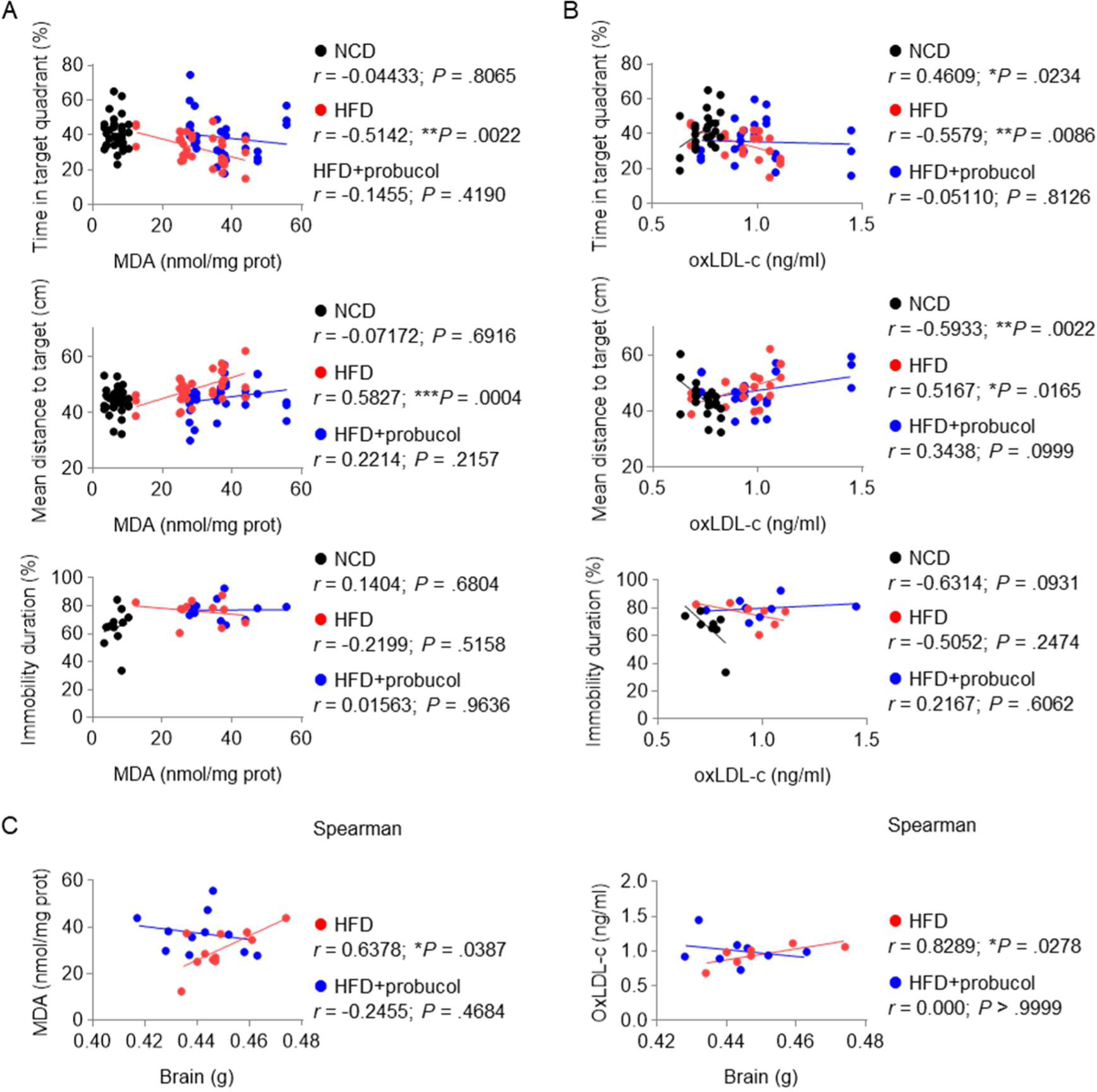
Probucol treatment dissociates MDA levels and cognitive decline in mice. (**A**) The correlations between MDA and behavior performance. The correlations are measured by Pearson r correlation. HFD-feeding started 8 weeks before treating the male C57BL/6J mice with 0.005% probucol in drinking water for 12 weeks. (**B**) The correlations between oxLDL-c and behavior performance. The correlations are measured by Pearson r correlation. (**C**) The correlations between MDA and the brain weight, and the correlation between oxLDL-c and the brain weight. The correlations are measured by Spearman r correlation.

### Probucol counteracts HFD’s impact by differentially regulating radical species

In order to examine the molecular mechanisms underlying the beneficial effects of probucol on cognitive performance, we analyzed the levels of candidate proteins known to mediate HFD-induced inflammation and oxidative stress in the brain. In the cortex, HFD-feeding resulted in a significant increase in the inducible form of nitric oxide synthase (iNOS), which is responsible for neurodegenerative changes in the cortex (68). Probucol treatment counteracted the effects of HFD on the induction of iNOS in the cortex, restoring its levels to that observed in the cortex of NCD-fed mice (Figure 6, A and B). However, in the hippocampus, while HFD had no effect on iNOS levels, probucol treatment increased iNOS levels (Figure 6, C and D). This is noteworthy as iNOS has been reported to play a role in adult neurogenesis (69).

**Figure 6.**
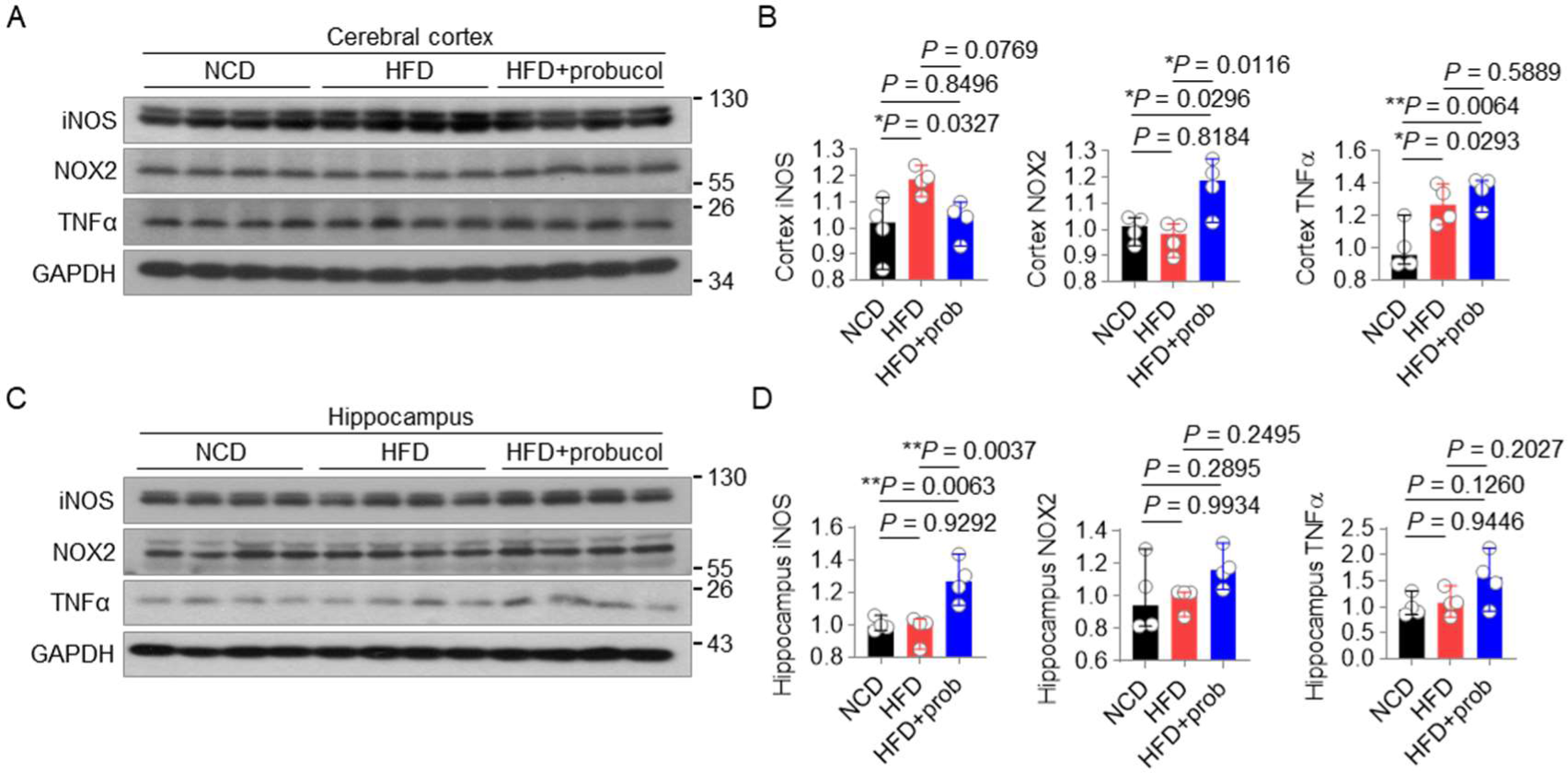
Probucol differentially regulates radical species in mice cerebral cortex. (**A-D**) Western blotting analysis of cerebral cortex and hippocampus lysates of three groups mice are shown in **A** and **C**. Brain extracts of male mice were analyzed by Western blotting using antibodies against iNOS, NOX2, TNFα and GAPDH. The levels of the proteins in the mice cortex and hippocampus are quantified in **B** and **D**, respectively (n = 4 mice per group, ordinary one-way ANOVA, followed by Tukey’s multiple comparisons test). HFD-feeding started 8 weeks before treating the male C57BL/6J mice with 0.005% probucol in drinking water for 12 weeks.

The levels of Nicotinamide Adenine Dinucleotide Phosphate oxidase 2 (NOX2), a major source of superoxide in the brain, were significantly increased in the cortex and moderately accumulated in the hippocampus of probucol-treated mice compared to both NCD-fed mice and untreated HFD-fed mice (Figure 6, A-D). Given the important roles of ROS in learning, memory, and brain plasticity (70, 71), it is possible that probucol combats the detrimental effects of HFD by upregulating NOX2. We also examined the levels of the inflammation marker TNFα. It was similarly upregulated in the both the untreated HFD-feeding mice and the probucol treated mice compared to the NCD-fed mice. However, there was no statistical difference in the levels of TNFα in the hippocampus between these groups (6, A-D). These data indicate that probucol differentially regulates the machineries for nitric oxide and superoxide without changing the inflammatory status in the cortex.

## Discussion

Fat-rich diet is regarded as an important factor not only for the development of metabolic disorders but also CNS abnormalities (72). In this study, the effects of probucol on HFD-fed mice are systematically examined in the aspects of cognition and mood-relevant behaviors, global metabolism and redox status. Probucol shows prominent benefits to antagonize the HFD-induced decline in spatial learning and memory, and improves sociability, while it has no effect on the anxiety or depression-like behaviors. Probucol treatment did not antagonize the metabolic alterations demonstrated by similarly elevated glucose, total cholesterol, LDL-cholesterol or oxidized LDL levels in the blood, which was consistent with previous studies performed in various animal models (56, 73, 74). Moreover, the levels of the oxidative stress marker MDA are unexpectedly increased together with a reduction of total GSH in the probucol treated mice. Investigation of the brain morphological and molecular changes demonstrated that the beneficial effects of probucol is associated with the restoration of cortex and hippocampus morphology, and differentially regulation of free radical production machineries.

By close measurement of the cortex and hippocampus, we found that HFD increased the area and width of the hippocampus, which is reduced probucol treatment. Of note, it has also been demonstrated that probucol treatment moderately reduced hippocampal dentate gyrus volume in FVB/N mice (75). Although reduction in hippocampus is frequently observed in human and mice models with cognitive decline, it is worth noting that increased hippocampus size is also associate with cognitive deficits such as decreased social interaction and delayed learning capacity in some mice models (76, 77). Consistently, the positive correlation between brain weight and body weight in untreated HFD-fed mice indicate that neuronal cell hypertrophy or hyperplasia in the hippocampus could be detrimental, and destruction of such correlation may contribute to an improved CNS function by probucol.

The relevance of oxidative stress in obesity-associated comorbidities has been intensely studied (78, 79). It is generally considered that overproduction of free radicals and related inflammatory conditions is detrimental, while these factors are also found essential for normal physiological response under specific stressed conditions such as cold-induced brown adipocyte thermogenesis (80). Contrary to the potential antioxidant properties of probucol demonstrated in severely diseased models, we found that systemic and regional oxidative stress seem to be even increased in probucol-treated mice that are fed with HFD. Similar to this data, probucol was also reported to upregulate erythrocyte and plasma lipid peroxidation in mice and macaques (81, 82) and serum NO levels in Sprague Dawley rats (83), enhances NO bioactivity in aortic rings in rabbits (84). Considering the low LDL-c plasma lipoprotein distribution in mice compared with human (85) and the reliance of incorporation into the LDL particles for the antioxidant effect of probucol (), it remains to be decided whether the reported potential antioxidant effect of probucol reported in patients with very high LDL levels in the blood such as those with familial hypercholesterolemia is similarly reproduced in specific mice models. In addition, possibly due to the low solubility and cell permeability, probucol has no effect on liver weight or liver cholesterol (53). Beta-carotene that has no effect on LDL oxidation can effectively prevent lesion formation to the same degree as probucol in cholesterol fed rabbit (86). Moreover, the metabolic/antioxidant function and the CNS effects of probucol are not consistently observed in animal models (56, 73).

Taken together, the current study demonstrates the potential effect of probucol in antagonizing HFD-induced cognitive decline without bringing systemic metabolic benefits and reduction of oxidative stresses. These findings would prompt a rethinking of the functioning mechanism of probucol as well as the roles of altered metabolic profiles and free radicals in brain function. These findings suggest that it is important to re-examine the roles of metabolic shift, redox homeostasis and inflammation in the development of diet-induced cognitive deficits.

## Methods

### Morris navigation swim test

Morris navigation swim task was performed as previously described with some modifications (87, 88). The water maze comprised of a round tank with a diameter of 90 cm, filled with water maintained at approximately 22℃ (Xiamen Baocheng Biotechnology, Xiamen, China). In each quadrant of the pool wall, graphic clues were affixed. The tasks were conducted from 6:00 pm to 10:00 pm. Throughout the navigation training, a fixed platform was placed in one of the four quadrants. The mice underwent training for 7 days, with four trials per day, starting from four different locations in a pseudorandom manner. Training continued until 90% of the mice successfully located the platform within 30 s for two consecutive days. On the eighth day, spatial memory capacity was evaluated using a spatial probe test, wherein the platform was removed. The mice were given 60 s to search for the original location of the platform. The performance of the mice was recorded and analyzed using an automatic tracking system (SMARTPREMIUM 3.3, Panlab Harvard Apparatus, Barcelona, Spain). The latency to target and the area under the curve (AUC) of latency to target during acquisition are analyzed to assess spatial learning and memory. The percentage of time spent in the target quadrant, mean distance to target, and total crossing numbers during the probe trials are analyzed to evaluate the spatial navigation abilities of the mice.

### Three-chamber social approach task

The three-chamber social approach task was conducted as previously described with some modifications (89). The apparatus used was a rectangular box divided into three chambers, measuring 60 cm × 40 cm (Xiamen Baocheng, Fujian, China). The test mice were placed in the middle chamber and given access to both end chambers, each equipped with a wired cage. During the habituation phase, the cages in both end chambers were empty. In the sociability test, one stranger mouse was placed inside one of the cages, while the other cage remained empty. In the social preference test, a second stranger mouse was placed inside the wired cage in the opposite side chamber. Each session lasted 5 minutes for the habituation phase, 10 minutes for the sociability test, and 5 minutes for the social preference test. The movement and interactions of the mice were recorded and analysed using an automatic tracking system (SMARTPREMIUM 3.3, Panlab Harvard Apparatus, Barcelona, Spain). The interaction region was defined as a 3 cm area surrounding the wire cage.

### Elevated plus maze test

The elevated plus maze used in this study was a customized four-armed apparatus. Each arm measured 30.8 cm × 6 cm × 16 cm (Xiamen Baocheng, Fujian, China) and was elevated 65 cm off the floor. The mice were placed in the center of the maze, facing the closed arms, and allowed to explore freely for a duration of 5 minutes. The movement and behavior of the mice were recorded and analyzed by an automatic tracking system (SMARTPREMIUM 3.3, Panlab Harvard Apparatus, Barcelona, Spain). Additionally, the number of times of the mice explored downward was manually counted.

### Forced swim test

The test utilized a cylindrical water tank with a height of 30 cm and a diameter of 15 cm (Xiamen Baocheng, Fujian, China). The water level in the tank was maintained at approximately 15 cm above the bottom and kept at a temperature of around 22°C. The mice were released into the tank and allowed for freely explore for a duration of 6 minutes. The movements of the mice was recorded and analysed by an automatic tracking system (SMARTPREMIUM 3.3, Panlab Harvard Apparatus, Barcelona, Spain).

### Y maze

The Y maze used in this study is a customized apparatus with three 37.8 cm × 6.7 cm × 16 cm arms (Xiamen Baocheng, Fujian, China). The mice were placed in the center of the Y maze, facing the direction of one of the arms. They were allowed for freely explore the maze for a total of 8 minutes. The movements of the mice were recorded and analyzed using an automatic tracking system (SMARTPREMIUM 3.3, Panlab Harvard Apparatus, Barcelona, Spain).

### Immunoblotting

The left cerebral hemisphere of euthanized mice was isolated and the cerebral cortex and hippocampus were collected in ice-cold PBS. The tissues were homogenized and sonicated in RIPA buffer, which consisted of 1×PBS, 1% NP-40, 0.1% SDS, 0.6% sodium deoxycholate, and phosphatase and protease inhibitor cocktails. The homogenates were then centrifuged at 20000 *g* for 15 minutes. The resulting supernatants were collected and mixed with SDS-PAGE sample buffer. The mixture was heated at 70℃ for 15 minutes. Subsequently, the protein extracts were subjected to SDS-PAGE and electrophoretic transfer. Immunoblots were performed following the protocols provided by the manufacturers of the primary antibodies. In this study, the following primary antibodies were used for immunoblotting: iNOS Polyclonal Antibody (Proteintech, 18985-1-AP), NOX2 Polyclonal Antibody (Proteintech, 19013-1-AP), TNF Alpha Polyclonal Antibody (Proteintech, 17590-1-AP), and GAPDH antibody (Proteintech, 60004-1-Ig).

### Histological analysis

The mice brains were perfused with PBS, followed by fixation in 4% paraformaldehyde for 24 hours. The brains were then processed for OCT embedding and serially cut into sections measuring 10 µm using a Leica CM1950 freezing microtome. Each section was allowed to dry for 30 minutes at 56℃ and then washed three times in PBS for 5 minutes each. Subsequently, the sections were stained with hematoxylin and eosin for a duration of 20 seconds.

### Serum lipoprotein determination

Blood samples were collected and centrifugation at 4℃, 800 *g* for 15 minutes. The levels of serum total cholesterol (TC) and low-density lipoprotein cholesterol (LDL) were measured using a Chemistry Analyzer (BS-240vet, Mindray Bio-Medical Electronics, Shenzhen, China). To detect serum oxidized low-density lipoprotein (oxLDL), a mouse oxLDL ELISA kit (EM0400, Wuhan Fine Biotech, Wuhan, China) was used following the manufacturer’s instructions.

### MDA detection

The livers were isolated from the euthanized mice using ice-cold PBS. The tissues were homogenized and sonicated in a cold saline solution, followed by centrifugation at 2500 rpm for 15 minutes. The resulting supernatants were collected and mixed with the medium provided by MDA kit (A003-1, Nanjing jiancheng Bioengineering Institute, China). The mixtures were then heated at 95℃ for 50 minutes and subsequently centrifuged at 3500 rpm for 10 minutes. The supernatant was measured at 532 nm using a spectrophotometer, following the manufacturer’s instructions. The levels of MDA were expressed as the nanomoles per gram of protein sample.

### GSH detection

Liver samples were collected and homogenized in the medium provided by T-GSH/GGSH kit (A061-1, Nanjing jiancheng Bioengineering Institute, China). The homogenized samples were then centrifuged at 3500 rpm for 15 minutes. The resulting supernatant was measured at 405 nm using a spectrophotometer, following the manufacturer’s instruction. The levels of total glutathione (T-GSH) and reduced glutathione (GSSH) were expressed as micromoles per gram of protein sample.

### Statistical analyses

Statistical analyses were conducted using Prism software (GraphPad Software, La Jolla, CA, USA). For normally distributed data, significance between two groups was determined using an unpaired two-tailed Student’s *t* test. For non-normally distributed data, significance between the two groups was determined using an unpaired two-tailed Mann-Whitney test. For comparisons among multiple groups with two fixed factors, ordinary two-way independent measure ANOVA, or two-way repeated measure (RM) ANOVA was performed, followed by Holm-Sidak’s multiple comparisons test. Geisser-Greenhouse correction was used where applicable. For all data, differences were considered significant when *P* < 0.05. Statistical details can be found in the figure captions.

## Author contributions

Conceptualization, Yi-Hong Zhan and Shu-Yong Lin; Data curation, Han-Ming Wu, Na-Jun Huang and Shu-Yong Lin; Formal analysis, Han-Ming Wu, Na-Jun Huang, Yang Yang, Yue Xu, Jian-Zhen Chen and Shu-Yong Lin; Funding acquisition, Zheng-Hao Yao, Sheng-Cai Lin, Yi-Hong Zhan and Shu-Yong Lin; Investigation, Han-Ming Wu, Na-Jun Huang, Yang Yang, Li-Ping Fan, Tian-Yu Tang, Lin Liu, Ze-Xin Cai, Xin-Yi Ren and Zheng-Hao Yao; Methodology, Han-Ming Wu, Na-Jun Huang, Li-Ping Fan, Dong-Tai Liu, Xi Huang, Cixiong Zhang, Xiang You and Ying He; Project administration, Zhi-Yun Ye, Wei Hong, Yi-Hong Zhan and Shu-Yong Lin; Resources, Li-Ping Fan, Chen Wang, Ying He and Yi-Hong Zhan; Supervision, Yi-Hong Zhan and Shu-Yong Lin; Validation, Han-Ming Wu; Visualization, Han-Ming Wu and Shu-Yong Lin; Writing – original draft, Han-Ming Wu and Shu-Yong Lin; Writing – review & editing, Sheng-Cai Lin, Yi-Hong Zhan and Shu-Yong Lin.

## Acknowledgement

We thank Dr. Meng-Xi Niu all the other members of SCL laboratory for technical assistance. This work was supported by grants from the National Key Research and Development Project of China (#2022YFA0806500), National Natural Science Foundation of China (#31822027, #82088102), the Fundamental Research Funds for the Central Universities (#20720210110), the Natural Science Foundation of Fujian Province of China (2021J011356), the Science and Technology Program of Xiamen (3502Z20224ZD1006) and XMU Training Program of Innovation and Entrepreneurship for Undergraduates (#2020Y1023).

## Supplementary Figure

**Figure S1.**
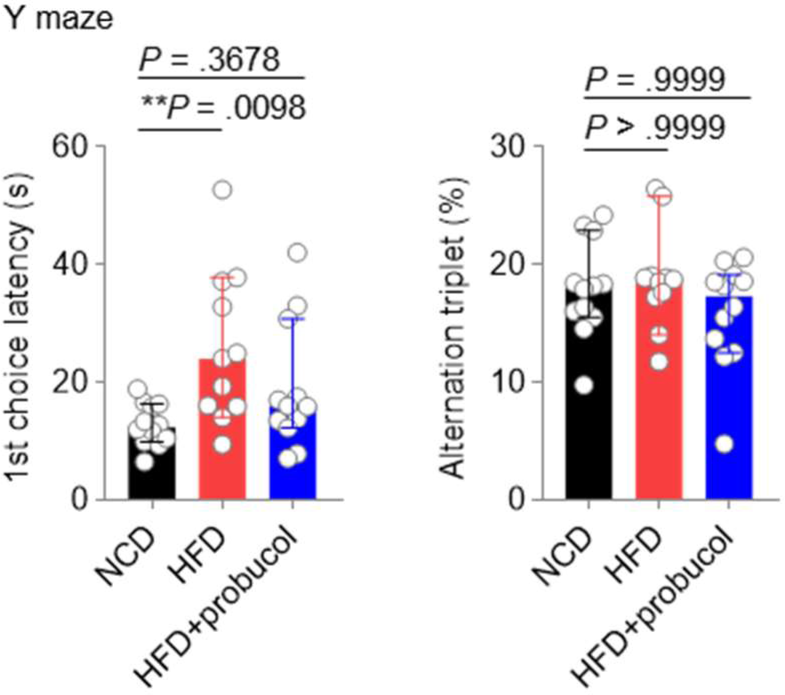
Probucol has no effect on the Y maze performance of the HFD-fed mice. HFD-feeding started 8 weeks before treating the male C57BL/6J mice with 0.005% probucol in drinking water for 12 weeks. The data of first choice latency and the percentage of alternation triplet are shown as individual values with median ± 95% CI (n = 11 or 12 mice for each group, Kruskal-Wallis test followed by Dunn’s multiple comparisons test).

## Notes

**Conflict of interest**: The authors have declared that no conflict of interest exists.

### Competing Interest Statement

The authors have declared no competing interest.

